# Oncomodulin (OCM) uniquely regulates calcium signaling in neonatal cochlear outer hair cells

**DOI:** 10.1101/2022.03.03.482327

**Authors:** Kaitlin E. Murtha, Yang Yang, Federico Ceriani, Jing-Yi Jeng, Leslie K. Climer, Forrest Jones, Jack Charles, Sai K. Devana, Aubrey J. Hornak, Walter Marcotti, Dwayne D. Simmons

**Author notes:** These authors contributed equally to this work (Kaitlin E. Murtha, Yang Yang).

## Abstract

In cochlear outer hair cells (OHCs), a network of Ca^2+^ channels, pumps and Ca^2+^-binding proteins (CaBPs) regulates the localization, spread, and magnitude of free Ca^2+^ ions. During early postnatal development, OHCs express three prominent mobile EF-hand CaBPs: oncomodulin (OCM), α-parvalbumin (APV) and sorcin. We have previously shown that deletion of *Ocm* (*Ocm*^-/-^) gives rise to progressive cochlear dysfunction in young adult mice. Here, we show that changes in Ca^2+^ signaling begin early in postnatal development of *Ocm*^-/-^ mice. While mutant OHCs exhibit normal electrophysiological profiles compared to controls, their intracellular Ca^2+^ signaling is altered. The onset of OCM expression at postnatal day 3 (P3) causes a developmental change in KCl-induced Ca^2+^ transients in OHCs and leads to slower KCl-induced Ca^2+^ transients than those elicited in cells from *Ocm*^-/-^ littermates. We compared OCM buffering kinetics with other CaBPs in animal models and cultured cells. In a double knockout of *Ocm* and *Apv* (*Ocm*^*-/-*^*;Apv*^*-/-*^), mutant OHCs show even faster Ca^2+^ kinetics, suggesting that APV may also contribute to early postnatal Ca^2+^ signaling. In transfected HEK293T cells, OCM slows Ca^2+^ kinetics more so than either APV or sorcin. We conclude that OCM controls the intracellular Ca^2+^ environment by lowering the amount of freely available [Ca^2+^]_i_ in OHCs and in transfected HEK293T cells. We propose that OCM plays an important role in shaping the development of early OHC Ca^2+^ signals through its inimitable Ca^2+^ buffering capacity.

## 1. Introduction

Outer hair cells (OHCs) are unique motor-sensory cells responsible for cochlear amplification and give rise to the exquisite sensitivity and frequency selectivity of mammalian hearing. Mammalian OHCs use a variety of organelles and proteins to regulate Ca^2+^ signaling. In OHCs, the influx of Ca^2+^ primarily comes through mechanoelectrical transduction (MET) channels located in the stereocilia and through voltage-gated Ca^2+^ channels localized at the basolateral membrane [1-3]. Calcium extrusion is accomplished via plasma membrane Ca^2+^-ATPase (PMCA2) pumps localized to OHC stereocilia [4, 5]. Within OHCs, Ca^2+^ is stored in an elaborate network of subsurface cisternae, which functions as a modified endoplasmic reticulum, as well as in mitochondria and the nucleus [6]. Additional key players controlling OHC Ca^2+^ dynamics are the developmentally regulated EF-hand Ca^2+^ binding proteins (CaBPs) that serve as mobile buffers [7, 8]. These components are all integral to the Ca^2+^ signaling network within OHCs and, prior to hearing onset, will undergo developmental changes important to the functional maturation of OHCs (**Figure 1A**).

**Figure 1.**
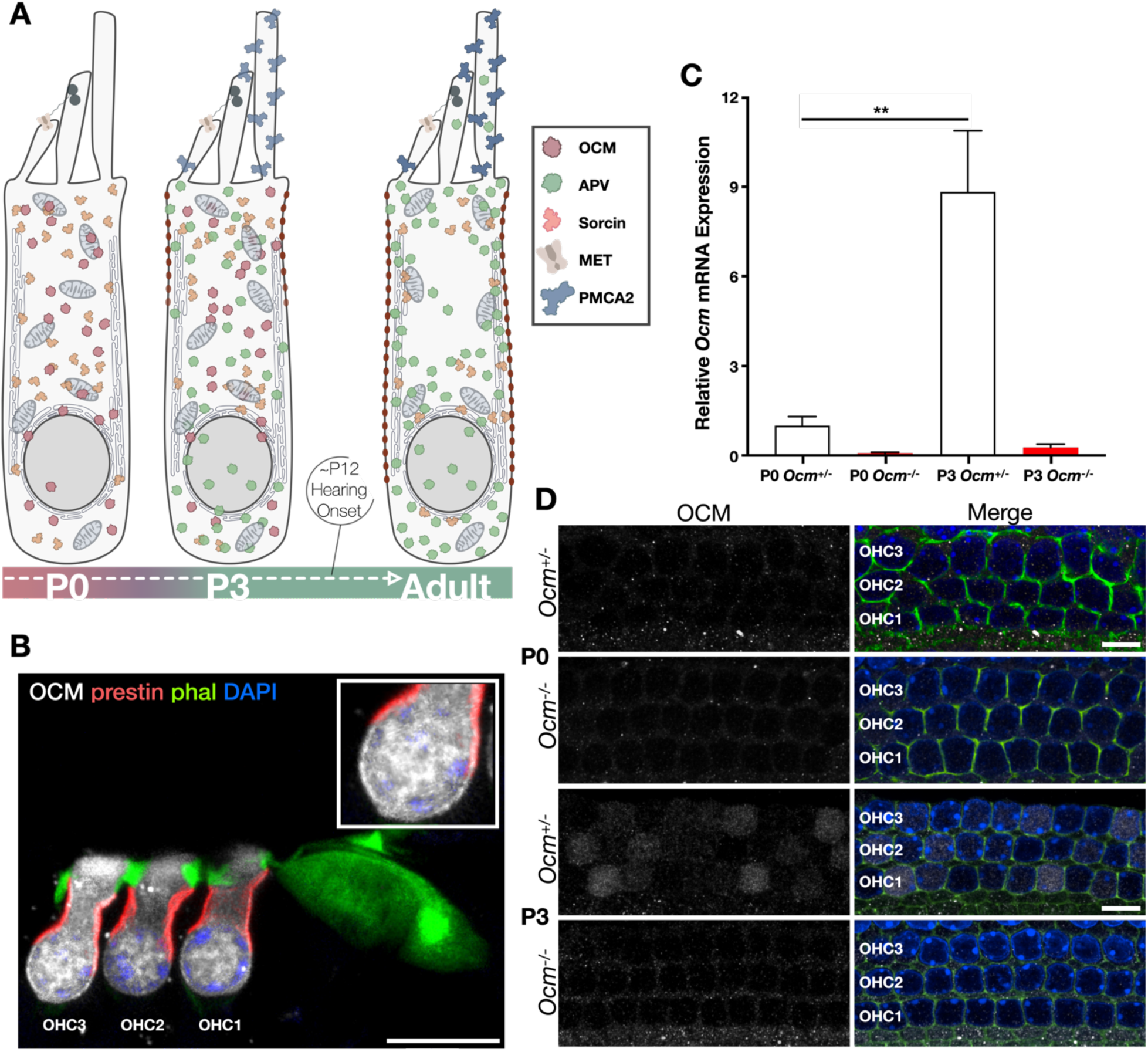
Expression of OCM begins early in development in mouse OHCs. **A)** Graphical timeline illustrating changes in the expression of Ca^2+^ related genes during OHC development (key on right). At P0, APV (red) expression is abundant [38]. Sorcin (orange) expression is also observed. By P3, OCM (green) protein expression is upregulated, while APV expression is downregulated [38]. **B)** Maximum Intensity Projection of mid-modiolar organ of Corti preparation from adult mouse (3 mo.). Sample was stained with OCM (white), prestin (red), phalloidin (green) and DAPI (blue). Inset shows details of OHC sub-nuclear region. **C)** Relative *Ocm* transcript levels measured via qRT-PCR in P0 and P3 *Ocm*^+/-^ and *Ocm*^-/-^ mice. Bars represent mean transcript levels (normalized to *B2M*, a housekeeping gene) and are plotted relative to P0 *Ocm*^+/-^. Error bars represent S.E.M. Significance (*t* test) is denoted as: *\*\*p=0.005*. **D)** Confocal images of cochlear spirals harvested from P0 and P3 *Ocm*^+/-^ and *Ocm*^-/-^ mice. OCM (white), phalloidin (green) and DAPI (blue) are shown. Scale bar = 10μm. Maximum Intensity Projections are composed of 2 × 1 μm slices. 63x magnification.

A wide variety of EF-hand CaBPs have been developmentally characterized in the cochlea [8-10], but their role in OHC maturation remains obscure. Developmental expression patterns of two parvalbumin isoforms, α-parvalbumin (*Apv*) and β-parvalbumin/oncomodulin (*Ocm*), are well documented in the organ of Corti [11-13]. At P0, APV protein expression is abundant and diffuse (**Figure 1A**) [12]. During OHC maturation, OCM protein expression increases dramatically, while APV expression is downregulated [12, 14, 15]. Recently, sorcin (*Sri*), a CaBP involved in maintaining ER Ca^2+^ homeostasis, was identified as a top cell-type defining gene for OHCs, along with *Slc26a5* (encoding the motor protein prestin) and *Ocm* [16, 17].

The most abundant CaBP in adult OHCs is OCM [12]. Following the onset of hearing in rodents, OCM localizes to the lateral membrane but is also abundant in the cytoplasm and nucleus (**Figure 1A,B**). Targeted deletion of *Ocm* leads to an early-onset hearing loss phenotype in both C57Bl/6 and CBA/CaJ mice [10, 18]. In both strains, hearing across lifetime (hearing span) is reduced by more than 50% in *Ocm* knockout mice compared to wildtype [18-20]. There is no known hearing loss phenotype associated with targeted deletion of any other CaBP [20, 21]. Downregulation of other CaBPs in favor of OCM during development, along with the hearing loss phenotype of *Ocm*^-/-^ mice in the adult, leads us to hypothesize that OCM is compulsory for maintaining hearing function.

Here, we explore OCM-dependent Ca^2+^ dynamics in developing OHCs from *ex vivo* cochlear explants and in a transfected mammalian cell line. We find that the absence of OCM leads to major changes in OHC Ca^2+^ signaling. These changes in Ca^2+^ signaling cannot be attributed to differences in the biophysical profiles of *Ocm*^*-/-*^ and *Ocm*^*+/-*^ OHCs. Although there are changes in APV and sorcin expression in response to loss of OCM, we conclude that they cannot compensate for OCM function. Experiments in cultured mammalian cells show that buffering by OCM differs from other Ca^2+^ buffers. Our results support the idea that OCM plays a unique and indispensable role as a Ca^2+^ buffer in developing OHCs.

## 2. Materials and methods

### 2.1. Animals

The original *Ocm* mutant mouse (C57Bl/6 *Actb*^*Cre*^*;Ocm*^*flox/flox*^) [18] was backcrossed onto the CBA/CaJ and the CBA/CaH background to minimize or eliminate any confounding effects of *Cdh23*^*ahl*^ mutation, which is linked to hearing loss in the adult [10]. Confirmation of congenicity was done by whole genome scan (Jackson Laboratories, Bar Harbor, ME). To generate *Ocm*^*-/-*^*;Apv*^*-/-*^ double mutants, *Ocm*^*-/-*^ mice were crossed with *B6.129P2-Pvalbtm1Swal/J* on the C57Bl/6 background (*Cdh23*^*ahl*^ mutation present) (Jackson Laboratories, Bar Harbor, ME). Animals were bred at the Baylor University Vivarium and at the animal facility of the University of Sheffield. Procedures performed in the UK and licensed by the Home Office under the Animals (Scientific Procedures) Act 1986 were approved by the University of Sheffield Ethical Review Committee. Experiments using CBA/CaH mice were performed at the University of Sheffield, where mice were sacrificed via cervical dislocation in accordance with UK Home Office regulations. All other experiments were performed at Baylor University and the University of California, Los Angeles (UCLA). Neonatal animals (P3 or less) were given a lethal injection of sodium pentobarbital (100-150 ng/ kg, IP) and immediately decapitated using scissors in accordance with US Public Health Service guidelines. For immunohistochemical analysis of adult mice, animals were given a lethal injection of sodium pentobarbital (100-150 ng/ kg, IP) and transcardially perfused. For hearing function analysis, Distortion Product Otoacoustic Emissions (DPOAEs) were performed as previously described [18]. All procedures performed in the USA were approved by the Baylor Institutional and UCLA Animal Care and Use Committees (IACUC) as established by the US Public Health Service and performed in compliance with the National Institutes of Health animal care guidelines.

### 2.2. Cloning

The rat *Ocm* gene (sequence in **Figure S01**) was cloned into pcDNA3.1CFP and pEGFP-N1 FLAG (Addgene #60360). Then, EGFP-FLAG was removed using BamHI (NEB) and NotI (NEB) and replaced with the mCherry sequence (gene block of mCherry from Qiagen) to produce p*Ocm*-mCh. Rat *Apv* and *Sri* (custom ordered gene block from IDT, Coralville, IA) (sequences in Figure **S01**) were swapped into the *Ocm* site using HindIII (NEB) /BamHI or HindIII/SalI to produce p*Apv-*mCh and p*Sri-*mCh, respectively. The mCherry plasmid also replaces the FLAG linker with a flexible GS linker (SGGGGSGGGGSGGGGS) between *Ocm;Apv;Sri* and mCherry. The mCh control plasmid was generated by cloning out the *Ocm* gene from *Ocm*-mCherry. Nucleotide sequences for rat *Ocm;Apv;Sri* and plasmid sequences for p*Ocm*-CFP, pmCh, p*Ocm*-mCh, p*Apv-*mCh and p*Sri-*mCh can be found in **Figure S01**.

### 2.3. Cell Culture and Reagents

Human Embryonic Kidney (HEK293T) cells were maintained in Dulbecco’s Modified Eagle Medium (DMEM, Thermo Fisher) supplemented with 10% fetal bovine serum (FBS, Bio-techne) at 37C in a 5% CO_2_ incubator. Transfections were performed using Lipofectamine 3000 (Thermo Fisher Scientific) in Opti-Mem reduced serum media (Thermo Fisher) according to manufacturer’s protocol. Mammalian cells were transfected with either mCherry (mCh), *Apv*-mCh, *Ocm*-mCh, *Sri*-mCh, or *Ocm*-CFP plasmids. 24 hours post transfection, growth media was replaced with HBSS (Gibco) before Ca^2+^ transient experiments.

### 2.4. Single-cell electrophysiology

OHCs from CBA/CaH *Ocm*^*+/-*^ and *Ocm*^*-/-*^ mice of both sexes were acutely dissected at P2 and P3. Cochleae were isolated from the inner ear as previously reported [22-25] using an extracellular solution composed of (in mM): 135 NaCl, 5.8 KCl, 1.3 CaCl_2_, 0.9 MgCl_2_, 0.7 NaH_2_PO_4_, 5.6 D-glucose, 10 HEPES-NaOH. Sodium pyruvate (2 mM), amino acids and vitamins were added from concentrates (Thermo Fisher Scientific, UK). The pH was adjusted to 7.5 (osmolality ∼308 mmol kg^-1^). The dissected cochleae were fixed at the bottom of the recording chamber by a nylon-meshed silver ring and perfused with the above extracellular solution. OHCs were viewed using an upright microscope (Olympus BX51) equipped with Nomarski Differential Interface Contrast (DIC) optics with a 60x water immersion objective and 15x eyepieces.

Recordings were performed at room temperature (21-24°C) using an Optopatch amplifier (Cairn Research Ltd, UK). Patch pipettes were pulled from soda glass capillaries and the shank of the electrode was coated with surf wax (2–3 MΩ). Current and voltage responses were measured using the following intracellular solution: 145 mM KCl, 3 mM MgCl_2_, 1 mM EGTA-KOH, mM 5 Na_2_ATP, 5 mM HEPES-KOH, 10 mM sodium phosphocreatine (pH adjusted to 7.28 with KOH; osmolality was 294 mmol kg^−1^). Voltage clamp protocols are referred to a holding potential of −84 mV. Data acquisition was performed using pClamp software (Axon Instruments, Union City, CA, USA) using a Digidata. Data were filtered at 2.5 kHz (8-pole Bessel), sampled at 5 kHz, and stored on computer. Offline data analysis was performed using Origin software (OriginLab, Northampton, MA, USA). Membrane potentials reported were corrected for the uncompensated residual series resistance (*R*_s_) and the liquid junction potential (LJP), which was 4 mV, measured between electrode and bath solutions.

### 2.5. Ca^2+^ imaging in neonatal OHCs

Three methods were used to measure Ca^2+^ dynamics in neonatal OHCs. For measurements using Ca^2+^ indicator dye Fluo-4, Ca^2+^ transients were induced either by local application of KCl or via KCl superfusion. For measurements using ratiometric Ca^2+^ indicator dye Fura-2, Ca^2+^ transients were induced via KCl superfusion. For all three methods, organ of Corti spirals were prepared as described previously [26, 27]. The apical third of the cochlea, corresponding to the 6-12 kHz region in the adult [28], was used.

For OHC Ca^2+^ transients induced by local application of KCl, cochlear spirals from P0 and P3 *Ocm*^*+/-*^ and *Ocm*^*-/-*^ CBA/CaH mice were harvested. Cochlear spirals were bathed in an extracellular solution composed of (in mM): 135 NaCl, 5.8 KCl, 1.3 CaCl_2_, 0.9 MgCl_2_, 0.7 NaH_2_PO_4_, 5.6 D-glucose, 10 HEPES-NaOH. Sodium pyruvate (2 mM), amino acids and vitamins were added from concentrates (Thermo Fisher Scientific, UK) [25, 29]. The pH was adjusted to 7.5 (osmolality ∼308 mmol kg^-1^). To access OHCs, Deiters’ cells were removed via suction of a small pipette (∼3-4 μm in diameter) as described previously [22, 25]. High K^+^ solution used for local application was (in mM): 113 NaCl, 1.3 CaCl, 0.9 MgCl_2_, 40 KCl, 10 HEPES, 5.6 Glucose, 0.7 NaH_2_PO_4_ (pH 7.5, osmolality ∼307 mmol kg^-1^). For Ca^2+^ dye loading, organ of Corti preparations were incubated for 40 min at 37°C in DMEM/F-12 (Gibco), supplemented with Fluo-4 AM (final concentration 10–20 µM; Thermo Fisher Scientific). The incubation medium contained pluronic F-127 (0.1% w/v, Sigma-Aldrich, UK) and sulfinpyrazone (250 µM) to prevent dye sequestration and secretion [27]. Preparations were transferred to a small microscope chamber, immobilized under nylon mesh, and brought to room temperature. To elicit Ca^2+^ transients, a ∼ 7 s local application of 40 mM KCl was delivered from a pipette positioned adjacent to row 3 OHCs. Ca^2+^ signals were recorded using a two-photon laser-scanning microscope (Bergamo II System B232, Thorlabs Inc., USA) based on a mode-locked laser system operating at 800 nm, 80-MHz pulse repetition rate and < 100-fs pulse width (Mai Tai HP DeepSee, Spectra-Physics, USA). Images were formed by a 60x objective, 1.1 NA (LUMFLN60XW, Olympus, Japan) using a GaAsP PMT (Hamamatsu) coupled with a 525/40 bandpass filter (FF02-525/40-25, Semrock). Using the two-photon microscope, fluorescence recording consisted of 4,000 frames taken at 30.3 frames per second from a 125 × 125 µm (512 × 512 pixels) region.

For OHC Ca^2+^ transients induced by KCl superfusion, cochlear spirals were isolated from P3 *Ocm*^*+/*^*;Apv*^*+/+*^, *Ocm*^*-/-*^*;Apv*^*+/+*^ or *Ocm*^*-/-*^*;Apv*^*-/-*^ mice (mixed CBA/CaJ and C57Bl/6 background), harvested, and prepared as described above. Cochlear spirals were bathed in a perilymph-like extracellular solution composed of (in mM): 136.8 NaCl, 5.8 KCl, 0.4 KH_2_PO_4_, 0.3 Na_2_HPO_4_, 0.8 MgSO_4_, 1.3 CaCl_2_, 4.2 NaHCO_3_, 5 HEPES and 5.6 glucose (pH 7.4-7.5, osmolality ∼306 mmol kg^-1^). Ca^2+^ signals were recorded using a custom-built confocal spinning disk (X-Light SDC with 6 laser lines, and a Photometrics sCMOS cooled camera) upright microscope (Leica, DM 6000 FS, Germany) with a 63x / 0.90 water immersion long working distance objective (Leica, 15506362 HCX APO L 63x/0.90 W U-V-I CS2). Using the spinning disc confocal microscope, each Ca^2+^ fluorescence recording includes 1500 frames taken at 80 frames per second using VisiView (VISITRON). KCl-induced depolarization solution was applied using a Picospritzer for bath perfusion. The solution was composed of (in mM): 142.4 KCl, 0.4 KH_2_PO_4_, 0.3 Na_2_HPO_4_, 0.8 MgSO_4_, 1.3 CaCl_2_, 4.2 NaHCO_3_, 5 HEPES and 5.6 glucose (pH 7.4-7.5, osmolality ∼305 mmol kg^-1^). Pressure was kept at a minimum (< 3 psi) to avoid triggering mechanically induced Ca^2+^ signals.

To measure intracellular Ca^2+^ concentration using the ratiometric dye, Fura-2, an upright Leica microscope (used above) was attached to a Lambda 10-B optical filter changer (Sutter instrument) with Lambda LS stand-alone xenon arc lamp (Sutter instrument), connected with pE-universal collimator (CoolLED). Fura-2 (final concentration 10 µM, Invitrogen) was administered to dissected cochlear spirals bathed in extracellular medium containing 0.2% F-127 (Invitrogen, USA). After 40 min incubation at 37 °C, the cochlear spiral was transferred to the microscope chamber and washed with extracellular solution to remove dye from the cell membrane before imaging. Spirals were depolarized using 37 mM KCl administered via superfusion. The intracellular free Ca^2+^ concentration was calculated using the emission intensity ratio from Fura-2 via follow this equation:

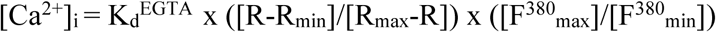

R is the ratio of 510 nm emission intensity with excitation at 340 nm to 510 nm emission intensity with excitation at 380 nm. R_min_ is the ratio at zero free Ca^2+^. R_max_ is the ratio at saturating Ca^2+^ (e.g., 39 μM). To measure the F^380^_max_ and F^380^_min_ in OHC, 25 μM ionomycin was applied to get the saturating intracellular Ca^2+^ (F^380^_max_), while F^380^_min_ in OHC was measured in Ca^2+^- free solution with 1mM EGTA when 25 μM ionomycin was applied. K_d_^EGTA^ was calculated and calibrated using Ca^2+^ calibration buffer kit (Life technologies, USA). A double log calibration plot of Fura-2 was calculated using the equation y = 0.5634x + 3.7127 with R^2^ = 0.9755. The Ca^2+^ response of the indicator is linear with the x-intercept being equal to the log of the apparent K_d_, 257.15 nM. The average of maximum Ca^2+^ concentration was calculated by the Fura-2 ratio of 510 nm emission intensity with excitation at 340 nm to 510 nm emission intensity with excitation at 380 nm.

Images were analyzed offline using custom-built software routines written in Python (Python 2.7, Python Software Foundation, RRID:SCR_014795) and ImageJ (NIH) [30]. Ca^2+^ signals were measured as relative changes of fluorescence emission intensity (*ΔF/F*_*0*_). *ΔF = ΔF - F*_*0*_, where *F* is fluorescence at time t and *F*_*0*_ is the fluorescence at the onset of the recording. After background subtraction, the activated OHCs were computed as pixel averages from square ROIs (size = 3.7 µm) centered on each OHC. Calcium transient rise-time constants were calculated using nonlinear regression in MATLAB using these equations (*x* represents time while *y* represents fluorescence intensity from each time point):

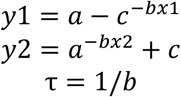

### 2.6. Ca^2+^ Transients of Transfected Mammalian Cells

HEK293T cells were incubated with Fluo-4 (10 µM, Invitrogen) for 30 minutes at 37°C. A puff of ionomycin (200 µL of 10 µM, Sigma) or ATP (100 µM, Sigma) was added to the perfusion chamber by a nearby pipette to induce Ca^2+^ transients. In a set of control samples, HEK293T cells were pre-incubated with cyclopiazonic acid (CPA, 10 µM), a SERCA inhibitor, to confirm that the source of Ca^2+^ was from ER stores [31]. A spinning disk confocal microscope was used to record changes in fluorescence over a 120 s time period at a frame rate of 6.25 frames/s. HEK293T cells were transfected with CFP plasmids and incubated with Fluo-4. To prevent spectral overlap between fluorophores, images were captured sequentially using excitation band filters of 440 -460 nm for CFP and 480 – 500 nm for Fluo-4 (**Figure S02**). A spinning disk confocal microscope was used to record changes in fluorescence over a 300 s time period at a frame rate of 1 frame/ 3 s for each filter. To categorize CFP expression levels, ROIs of individual cells were taken and fluorescence of cells (at least 100 for each group) were measured using ImageJ software. Data was transferred into MATLAB. Changes in fluorescence of single cells were plotted. Mean *ΔF/F*_*0*_ was calculated 30 seconds post-stimulation. Traces with obvious artifacts (e.g., due to motion) were removed from the analysis.

### 2.7. RT-PCR

*Ocm*-CFP transfected and non-transfected HEK293T cells were harvested and lysed using RLT buffer (Qiagen). RNA was isolated using the RNeasy plus mini kit (Qiagen). Reverse transcription reactions were performed using Transcriptor Reverse Transcriptase Kit (Roche). cDNA was subject to PCR using a Taq DNA Polymerase kit (Roche) using primers specific for mouse *Ocm* and *Glyceraldehyde 3-Phosphodehydrogenase (G3PDH)*. Primer sequences are provided in **Figure S03**. PCR cycling conditions were determined using the manufacturer’s protocol. A negative (no template) control was included. PCR amplicons were run on a 2% agarose EtBr gel and imaged.

### 2.8. qRT-PCR

Organ of Corti spirals were harvested from P0 and P3 CBA/CaJ mice, placed into RLT buffer (Qiagen), and immediately prepared for RNA extraction using the RNeasy Micro Kit (Qiagen) according to the manufacturer’s protocol. cDNA was synthesized using iScript Reverse Transcription Supermix (BioRad) according to manufacturer’s protocols. A BioRad CFX96 RT system was used to perform qRT-PCR using SsoAdvanced Universal SYBR Green (BioRad). n=3 technical replicates (i.e., 3 replicates for each sample per gene) were used for each plate. Expression levels for target genes were normalized to Beta-2-Microglobulin (*B2M*), a cochlear reference gene [32]. Primer sequences can be found in **Figure S03**. Quantification of mRNA expression (fold change) from the Cq data was calculated using the ΔΔCq method [33]. In short, the mRNA expression level of a target gene was first normalized to the average mRNA expression level of the corresponding reference gene (*B2M*) to obtain the ΔCq value. Then, the ΔΔCq of each gene was calculated using the following formula: ΔCq (target gene from sample) - ΔCq (target gene from P0 WT group). Then, 2^-ΔΔCq^ was calculated to represent the relative mRNA expression (fold change). All values were normalized to the mean Cq value of *Ocm*^+/-^ at P0. Raw qRT-PCR values and calculations can be found in **Figure S04**.

### 2.9. Immunofluorescence

Cochlea from neonatal CBA/CaJ mice (P0 and P3) were harvested and flushed with 4% paraformaldehyde (PFA, Sigma). Cochlea surface preparations were placed in sucrose for 30 minutes with shaking, fixed in 4% paraformaldehyde for 40 minutes, blocked with 5% NHST (0.3% Triton X-100) for 1 hour at room temperature. Samples were stained with antibodies to OCM (Santa Cruz sc-7446, 1:200), APV (SWANT PVG-213, 1:200), and sorcin (Invitrogen PA5-64975, 1:200). Primary antibodies were incubated overnight at 37C. Phalloidin-iFluor 488 (Abcam ab176753, 1:1000) and appropriate Alexa Fluor (Thermo 1:200) and Northern Lights (R&D Systems 1:200) conjugated secondary antibodies were incubated for 2 hours at 37ºC. Slides were prepared using Vectashield mounting media with DAPI (Vector Labs).

### 2.10. DPOAE Measurements

Hearing function was measured via distortion product otoacoustic emissions (DPOAEs) as described previously [18, 34, 35]. 1 month-old *Ocm*^*+/+*^*;Apv*^*+/+*^, *Ocm*^*-/-*^*;Apv*^*+/+*^, and *Ocm*^*-/-*^*;Apv*^*-/-*^ mice on C57Bl/6 were used. Acoustic stimuli were delivered using a custom acoustic assembly previously described [36]. Sound level was raised in 10 dB steps from 10 dB below threshold up to 80 dB sound pressure level (SPL). The frequencies that were used are as follows: 5.66, 8.00, 11.32, 16.00, 32.00, and 45.26 kHz. The Eaton-Peabody Laboratories Cochlear Function Test Suite (EPL-CFTS) was used to collect DPOAE input-output data and determine thresholds (f2 level required to produce a DPOAE at 0 dB SPL).

### 2.11. Statistical analysis

Statistical analysis was performed using GraphPad Prism 8 software. To compare means between two groups, an unpaired *t* test was used. For data sets that were not normally distributed, a Mann-Whitney U test was performed. When data was normally distributed, but standard deviations (SD) differed between groups, a Welch’s *t* test was used. To compare the means of three or more groups, an ANOVA was used. When normal distribution could not be assumed, a nonparametric Kruskal-Wallis test, followed by a Dunn’s multiple comparisons test was performed. Animals of either sex were randomly assigned to the different experimental groups. No statistical methods were used to define sample size, which was selected based on previous published similar work from our laboratories. Animals were taken from multiple cages and breeding pairs unless otherwise indicated. The electrophysiological (but not imaging) experiments were performed blind to animal genotyping.

## 3. Results

### 3.1. Ocm expression in the neonatal cochlea begins at P0

The maturation of OHC function follows the expression of a variety of channels, pumps and buffers that modulate levels of Ca^2+^ (**Figure 1A**). Although OCM is found throughout the cytoplasm in mature OHCs, it shows strong immunoreactivity (ir) near the lateral membrane subsurface cisternae and overlaps extensively with prestin-ir (**Figure 1B**). However, prior to prestin expression and OHC maturation, OCM is diffusely located throughout the cytoplasm (**Figure 1A**). In previous studies, we detected *Ocm* mRNA expression from whole cochlea as early as P3 [14]. To determine more precisely the onset of *Ocm* expression in the organ of Corti, we performed qRT-PCR and immunofluorescence on microdissected organ of Corti spirals from P0 and P3 mice. Low levels of the *Ocm* mRNA transcript could be detected in *Ocm*^*+/-*^ mice as early as P0, which then increased by approximately 9-fold by P3 (**Figure 1C**). While OCM-ir was absent at P0, P3 OHCs from *Ocm*^*+/-*^ mice showed low levels of OCM-ir, which varied in expression between individual cells (**Figure 1D**). As expected, OHCs from *Ocm*^-/-^ mice had negligible amounts of *Ocm* mRNA expression and OCM-ir (**Figure 1C**,**D**). OCM expression is, therefore, substantially upregulated in the *Ocm*^*+/-*^ cochlea between P0 and P3, with diffuse and variable cytoplasmic distribution.

### 3.2. Onset of OCM changes Ca^2+^ signaling in OHCs

After determining the onset of OCM protein expression in OHCs (between P0 and P3), we investigated how OCM expression influences Ca^2+^ signaling during early postnatal development. To measure this, we administered Ca^2+^ indicator dye, Fluo-4, to dissected cochlear preparations. Deiters’ cells were removed to access OHCs directly. Then, Ca^2+^ transients were elicited via a local application of KCl from a pipette positioned adjacent to the 3^rd^ row of OHCs. The Ca^2+^-related change in fluorescence intensity was recorded as *ΔF/F*_*0*_.

At P0, OHCs showed spontaneous and rapid Ca^2+^ transients (**Figures S05, S06**). We have previously shown that these Ca^2+^ transients are abolished in Ca^2+^-free solution, indicating their dependence on extracellular Ca^2+^ [25]. In addition to spontaneous activity, all three rows of OHCs from both P0 *Ocm*^*+/-*^ and *Ocm*^*-/-*^ mice responded to KCl application (**Figure 2A**,**B, S05, S06**). There was little to no difference in responses between OHC rows (**Figure S04, S05**) (only responses from the third row of OHC are shown). At P0, OHCs from *Ocm*^*+/-*^ and *Ocm*^*-/-*^ mice showed similar responses to local application of KCl (P0 *Ocm*^*+/-*^ 9.44±0.72 vs P0 *Ocm*^*-/-*^ 9.48±0.38). At this postnatal age, there was no difference in rise-time constants (τ) between *Ocm*^*+/-*^ and *Ocm*^*-/-*^ OHCs (*t* test) (**Figure 2E**).

**Figure 2.**
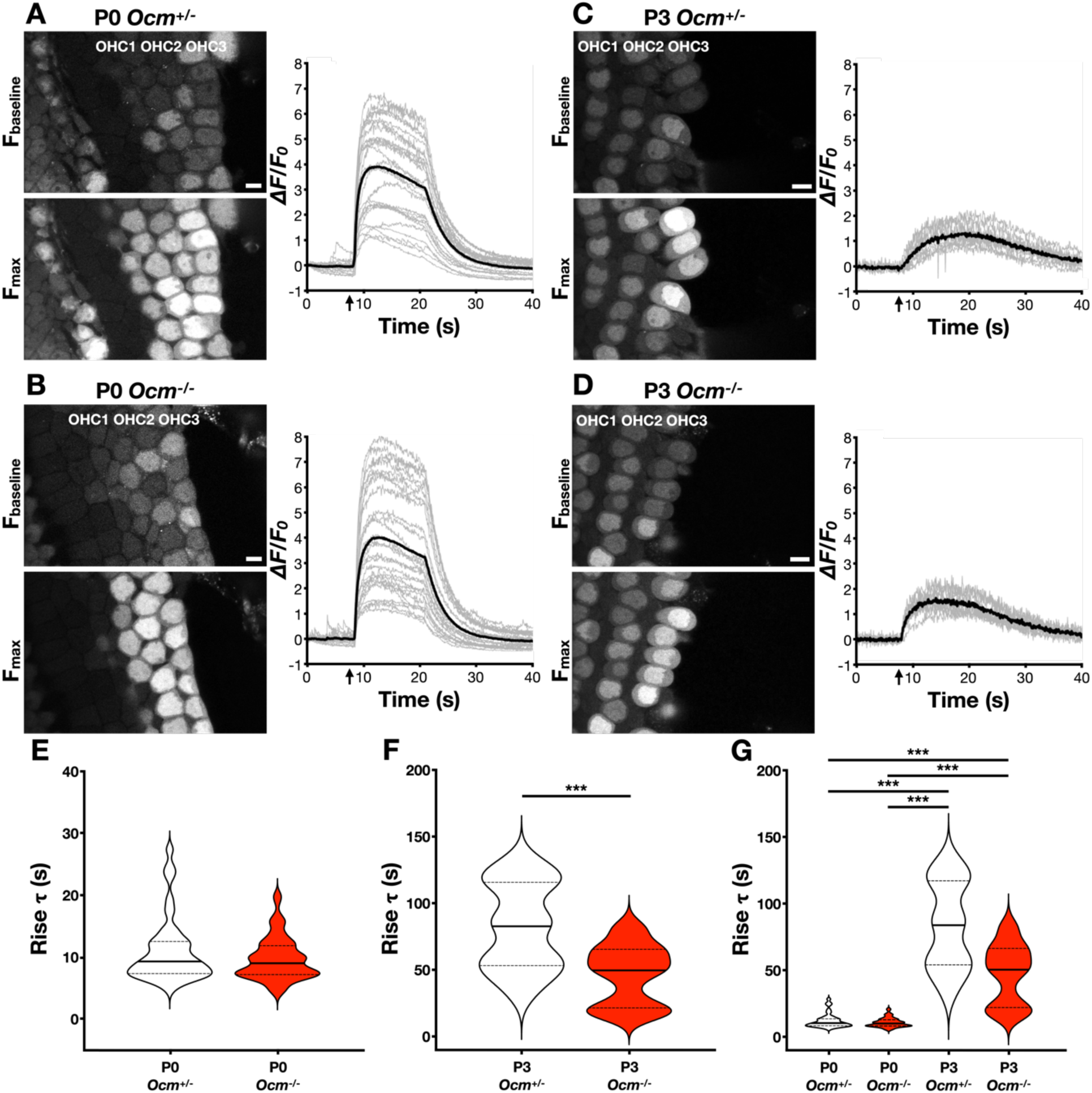
Calcium transients are induced by local application of KCl in P0 and P3 *Ocm*^+/-^ and *Ocm*^-/-^ OHCs. Acute organ of Corti preparations from P0 and P3 CBA/CaH mice were incubated with Fluo-4 and administered a local application of KCl. Representative images depict Fbaseline (before stimulation with KCl) and Fmax (maximum change in fluorescence). Scale bar = 10 μm. To the right are *ΔF/F0* plots for each genotype and age. Grey lines are *ΔF/F0* measurements from individual OHCs. Solid black lines represent the mean *ΔF/F0* for that preparation. Local application of KCl (arrow) was administered to **A)** P0 *Ocm*^+/-^ **B)** P0 *Ocm*^-/-^ **C)** P3 *Ocm*^+/-^ and **D)** P3 *Ocm*^-/-^ mice. Fluorescent transients were restricted to OHC3 (OHC row 3). **E-G)** Violin plots from KCl induced Ca^2+^ transient experiments in OHCs from P0 and P3 *Ocm*^+/-^ (white) and *Ocm*^-/-^ (red). Solid lines represent medians. Dotted lines represent quartiles. **E)** Rise τ from P0 *Ocm*^+/-^ (n=55 OHCs) and *Ocm*^-/-^ OHCs (n=55 OHCs). **F)** Rise τ from P3 *Ocm*^+/-^ (n=31 OHCs) and *Ocm*^-/-^ OHCs (n=21 OHCs) **G)** Rise τ compared between *Ocm*^+/-^ and *Ocm*^-/-^ P0 and P3 OHCs. Significant differences (*Mann-Whitney* test) denoted as follows: *\*\*\*p<0.0001*.

In contrast to P0, P3 OHCs exhibited less spontaneous Ca^2+^ transient activity (**Figure S07, S08**), which is consistent with other studies [25, 37]. The rise τ were different between P3 *Ocm*^*+/-*^ and *Ocm*^*-/-*^ OHCs (*t* test, *p<0.001*) (**Figure 2F**). P3 *Ocm*^+/-^ mice had a mean rise τ of 137.40±41.23, which was significantly slower compared to aged-matched *Ocm*^-/-^ OHCs that had a mean rise τ of 56.82±14.97 (**Figure 2F**). Regardless of genotype, the mean rise τ was considerably faster in P0 OHCs compared to P3 OHCs (*t* test, *p<0.001*) (**Figure 2G**).

### 3.3. Neonatal Ocm^-/-^ OHCs exhibit normal biophysical profiles

We then investigated whether differences in Ca^2+^ signaling in P3 OHCs between *Ocm*^+/-^ and *Ocm*^-/-^ mice (**Figure 3**) had any effect on their biophysical properties by performing patch-clamp experiments. Potassium currents, which at this age are primarily carried by delayed rectifier K^+^ channels [22], were recorded while applying a series of hyperpolarizing and depolarizing voltage steps (in 10 mV increments) from the holding potential of -84mV (**Figure 3A**,**B**). The time-course and voltage-dependence of the outward K^+^ currents in OHCs were comparable between *Ocm*^*+/-*^ and *Ocm*^-/-^ mice (**Figure 3A-C**). The total outward K^+^ current measured at 0 mV was comparable between the two genotypes (**Figure 3D)**. Voltage responses to current injections were also indistinguishable between the two genotypes, with action potentials being elicited by depolarizing current injections (**Figure 3E**,**F**), as previously shown [22]. The resting membrane potential of OHCs, which was obtained by the voltage-clamp recordings, was not significantly different between *Ocm*^*+/-*^ and *Ocm*^-/-^ mice (**Figure 3G**).

**Figure 3.**
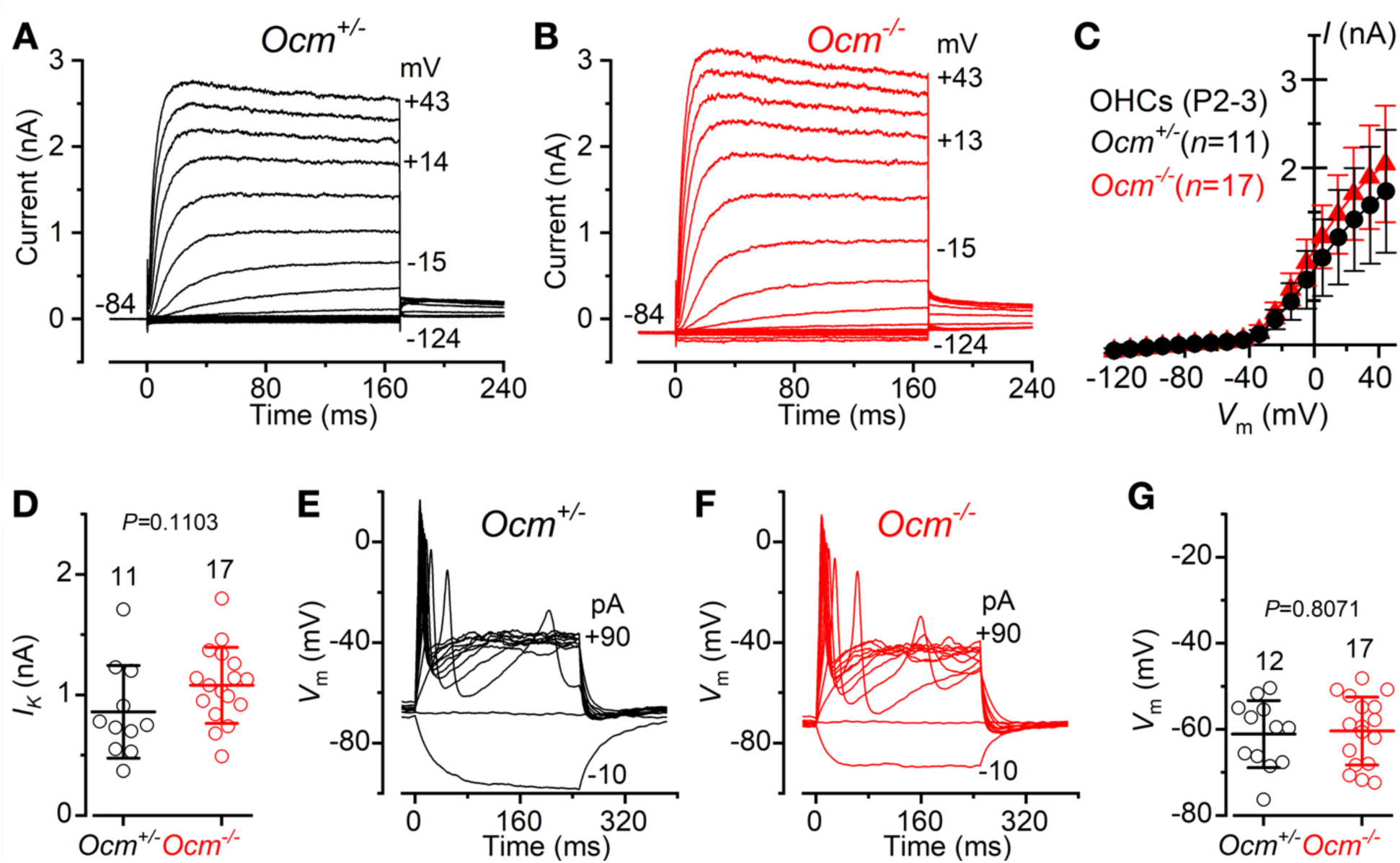
The biophysical properties of OHCs are not affected in early postnatal O*cm*^*-/-*^ mice. K^+^ currents from OHCs of P3 **A**) *Ocm*^*+/-*^ and **B**) *Ocm*^*-/-*^ CBA/CaH mice. Recordings were performed by applying a series of voltage steps in 10 mV nominal increments from the holding potential of -84 mV. **C)** Average current-voltage curves from OHCs of P2-P3 *Ocm*^*+/-*^ and *Ocm*^*-/-*^. **D)** The size of the total steady-state outward K^+^ current in OHCs, which was measured at 0 mV from the holding potential of -84 mV, was not significantly different between *Ocm*^*+/-*^ and *Ocm*^*-/-*^ (*p=0.1103, t* test). Voltage responses elicited by applying a series of 10 pA depolarizing current injections to OHCs of P3 *Ocm*^*+/-*^ **E**) and *Ocm*^*-/-*^ **F**) mice. **G)** Resting membrane potential recoded from OHCs of P2-P3 *Ocm*^*+/-*^ and *Ocm*^*-/-*^ mice. Values were not significantly different between the two genotypes (*p=0.8071, t* test). Single value recordings are shown as open symbols. The number of OHCs measured is shown above the average data points.

These experiments demonstrate that OHCs from *Ocm*^*-/-*^ mice have similar biophysical properties as those measured in control cells, indicating that the absence of OCM does not affect OHC electrophysiological function at these early postnatal ages.

### 3.4. Apv and Sri gene expression is altered in organ of Corti from Ocm^-/-^ mice

In some instances, the deletion of a particular CaBP is compensated for by a similar CaBP to maintain Ca^2+^ homeostasis [38-40]. For example, in the brain, OCM is upregulated in response to genetic deletion of *Apv* [39]. Thus, we investigated whether the loss of OCM alters the expression of two other CaBPs highly expressed in OHCs: APV and sorcin.

First, we quantified the relative mRNA expression levels of *Apv* and *Sri* in dissected organ of Corti spirals from P0 and P3 *Ocm*^*+/-*^ and *Ocm*^*-/-*^ mice. *Apv* transcript abundance was higher in P0 *Ocm*^*+/-*^ mice compared to age-matched *Ocm*^*-/-*^ mice, but this difference was not statistically significant (*t* test, two-tailed) (**Figure 4A**). At P3, the two genotypes had similar levels of *Apv* mRNA. *Sri* mRNA levels were comparable between P0 *Ocm*^*+/-*^ and *Ocm*^*-/-*^ mice (**Figure 4B**). At P3, *Ocm*^*-/-*^ mice had two-fold lower *Sri* mRNA expression compared to P3 *Ocm*^*+/-*^ (*t* test).

**Figure 4.**
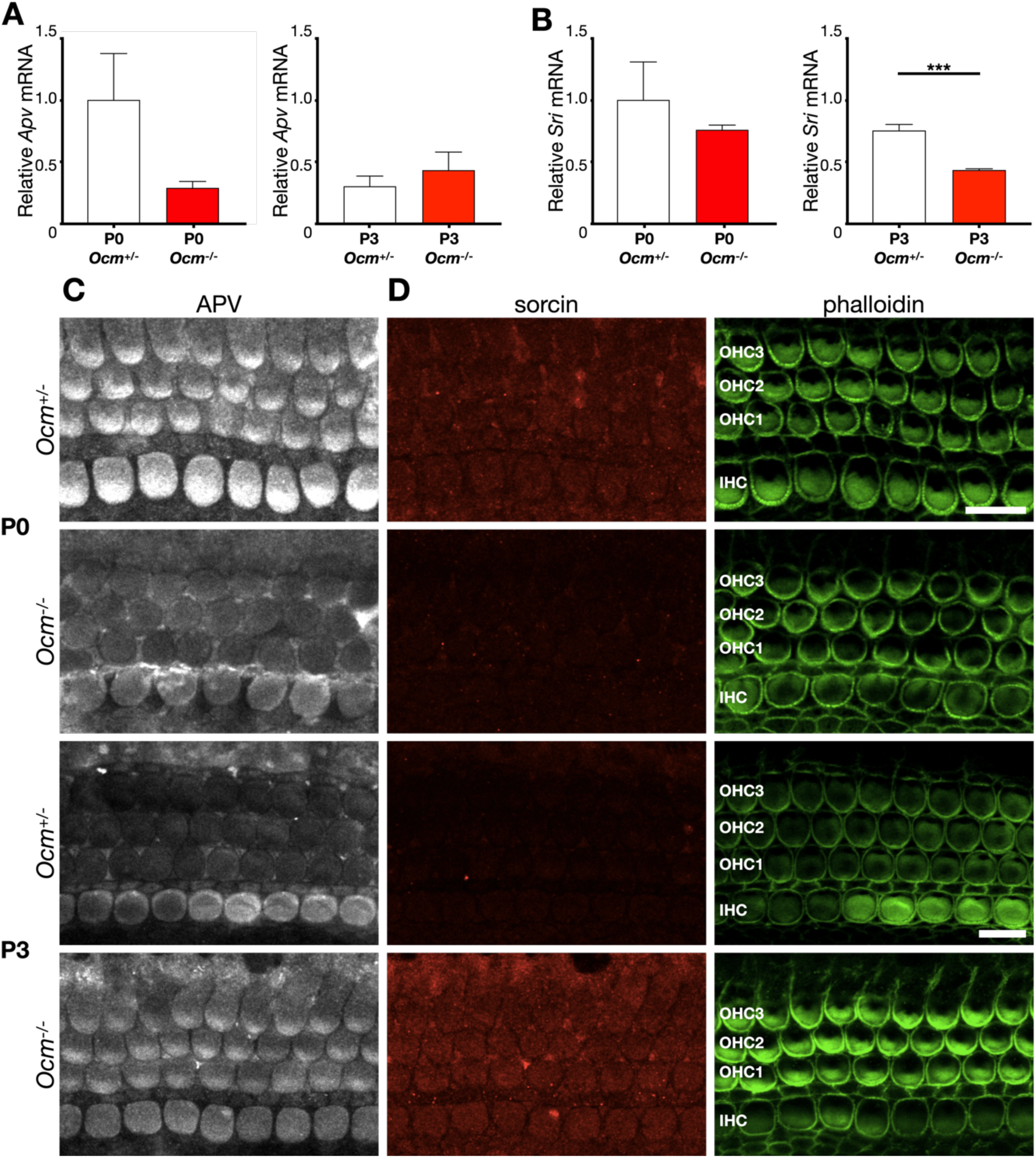
Absence of *Ocm* alters APV and sorcin protein expression levels. **A-B)** qRT-PCR was used to determine relative transcript levels in cochlear spirals harvested from P0 and P3 *Ocm*^+/-^ (white) and *Ocm*^-/-^ (red) mice. Mean values of relative normalized expression of **A)** *Apv* and **B)** *Sri* are represented by a bar graph. Mean transcript levels are plotted relative to P0 *Ocm*^+/-^ mean values. Error bars represent S.E.M. Significance (*t* test) is denoted as: *\*\*\*p<0.001*. Of note, when a one-tailed t test was used to compare mean values of *Apv* mRNA abundance between P0 *Ocm*^+/-^ vs *Ocm*^-/-^, there is a significant difference (*p=0.048*) **C-D)** Confocal images of cochlear spirals microdissected from P0 and P3 *Ocm*^+/-^ and *Ocm*^-/-^ mice. **C)** APV (white) and **D)** sorcin (red) were detected at P0 and P3 in both *Ocm*^+/-^ and *Ocm*^-/-^ mice. Phalloidin and DAPI (not shown) were used to outline OHCs and denote subcellular location. Maximum Intensity Projections of 1 μm slices starting from the cuticular plate and ending in the nucleus are shown here. Scale bar = 10 μm.

Using immunofluorescence, we investigated protein expression of APV and sorcin in organ of Corti spirals microdissected from P0 and P3 *Ocm*^*+/-*^ and *Ocm*^*-/-*^ mice. Generally, APV-ir was more abundant in IHCs compared to OHCs (**Figure 4C**). At P0, APV-ir is stronger in *Ocm*^*+/-*^ OHCs and IHCs compared to age-matched *Ocm*^*-/-*^ mice. However, at P3, APV protein expression consistently showed an opposite trend, where APV-ir was increased in OHCs of *Ocm*^*-/-*^ compared to OHCs from *Ocm*^*+/-*^ mice (**Figure 4D**). Next, we investigated the relative protein expression of sorcin. Overall, sorcin-ir was similar between IHCs and OHCs. At P0, less sorcin-ir was detected in *Ocm*^*-/-*^ OHCs and IHCs compared to *Ocm*^*+/-*^ (**Figure 4D**). At P3, *Ocm*^*-/-*^ OHCs and IHCs express more sorcin protein compared to their *Ocm*^*+/-*^ littermates (**Figure 4D**), which is opposite of what was observed for *Sri* mRNA expression at P3. Interestingly, while *Ocm* expression is minimal in IHCs [17], the absence of OCM gives rise to changes in APV and sorcin immunofluorescence in both OHCs and IHCs.

### 3.5. Loss of Apv further alters Ca^2+^ responses in OHCs from Ocm^-/-^ but does not affect hearing thresholds

Triple knockout of *Apv, calbindin-D28k*, and *calretinin* results in no observable hearing-related phenotype in 3–4-month-old mice [40]. *Ocm* is the only CaBP that when deleted, results in a hearing loss phenotype [10, 18]. However, up until the onset of hearing loss, *Ocm*^*-/-*^ OHCs appear functional and intact. We investigated whether the presence of APV was sufficient for maintaining OHC function in the absence of OCM. We measured the hearing thresholds of single *Ocm* knockout and double (*Ocm;Apv)* knockout mice. DPOAE thresholds are a standard way of assessing OHC function *in vivo*. We found that *Ocm*^*-/-*^*;Apv*^*-/-*^ mice did not have elevated hearing thresholds at 1-month of age compared to single *Ocm* knockouts (*Ocm*^*-/-*^*;Apv*^*+/+*^) (**Figure S09**). At 32 kHz, the *Ocm*^*-/-*^*;Apv*^*-/-*^ mice (44.03±12.59 dB SPL) displayed DPOAE thresholds that fell in between *Ocm*^*+/+*^*;Apv*^*+/+*^ (33.63±7.65 dB SPL) and *Ocm*^*-/-*^*;Apv*^*+/+*^ (54.14±12.28 dB SPL) (**Figure S09**). The only significant difference in DPOAE thresholds was between *Ocm*^*+/+*^*;Apv*^*+/+*^ and *Ocm*^*-/-*^*;Apv*^*+/+*^ mice (*p=0.0021*) (*one-way ANOVA*). Without expression of OCM or APV, the OHCs of young adult *Ocm*^*-/-*^*;Apv*^*-/-*^ mice remain functional and loss of APV does not further elevate hearing thresholds.

Since APV is expressed at significant levels during early OHC development, we investigated whether Ca^2+^ signaling would be further altered with the deletion of both *Ocm* and *Apv*, together. We recorded Ca^2+^ transients from OHCs in acutely dissected cochlear spirals isolated from P3 *Ocm*^*+/-*^*;Apv*^*+/+*^, *Ocm*^*-/-*^*;Apv*^*+/+*^, and *Ocm*^*-/-*^*;Apv*^*-/-*^ mice. Intact cochlear spirals required the use of KCl superfusion to elicit Ca^2+^ transients. OHCs from *Ocm*^+/-^*;Apv*^+/+^ mice had the lowest magnitude of response to KCl superfusion compared to either *Ocm*^-/-^*;Apv*^+/+^ or *Ocm*^-/-^*;Apv*^-/-^ (**Figure 5A, Figures S10-12**). To maintain consistency with previous Ca^2+^ transient experiments, the change in fluorescence intensity as *ΔF/F*_*0*_ resulting from KCl superfusion was recorded from row 3 OHCs only. As seen in **Figure 5A**, the *ΔF/F*_*0*_ responses were highest for OHCs from *Ocm*^-/-^*;Apv*^+/+^ and *Ocm*^-/-^*;Apv*^-/-^ mice. The rise-time constants of the KCl-induced Ca^2+^ signaling in OHCs from both *Ocm*^*-/-*^*;Apv*^*+/+*^ and *Ocm*^*-/-*^*;Apv*^*-/-*^ mice were significantly faster compared to control mice (*Ocm*^*+/-*^*;Apv*^*+/+*^) (*p=0.0459* and *p<0.0001*, respectively, *one-way ANOVA*, **Figure 5B**).

**Figure 5.**
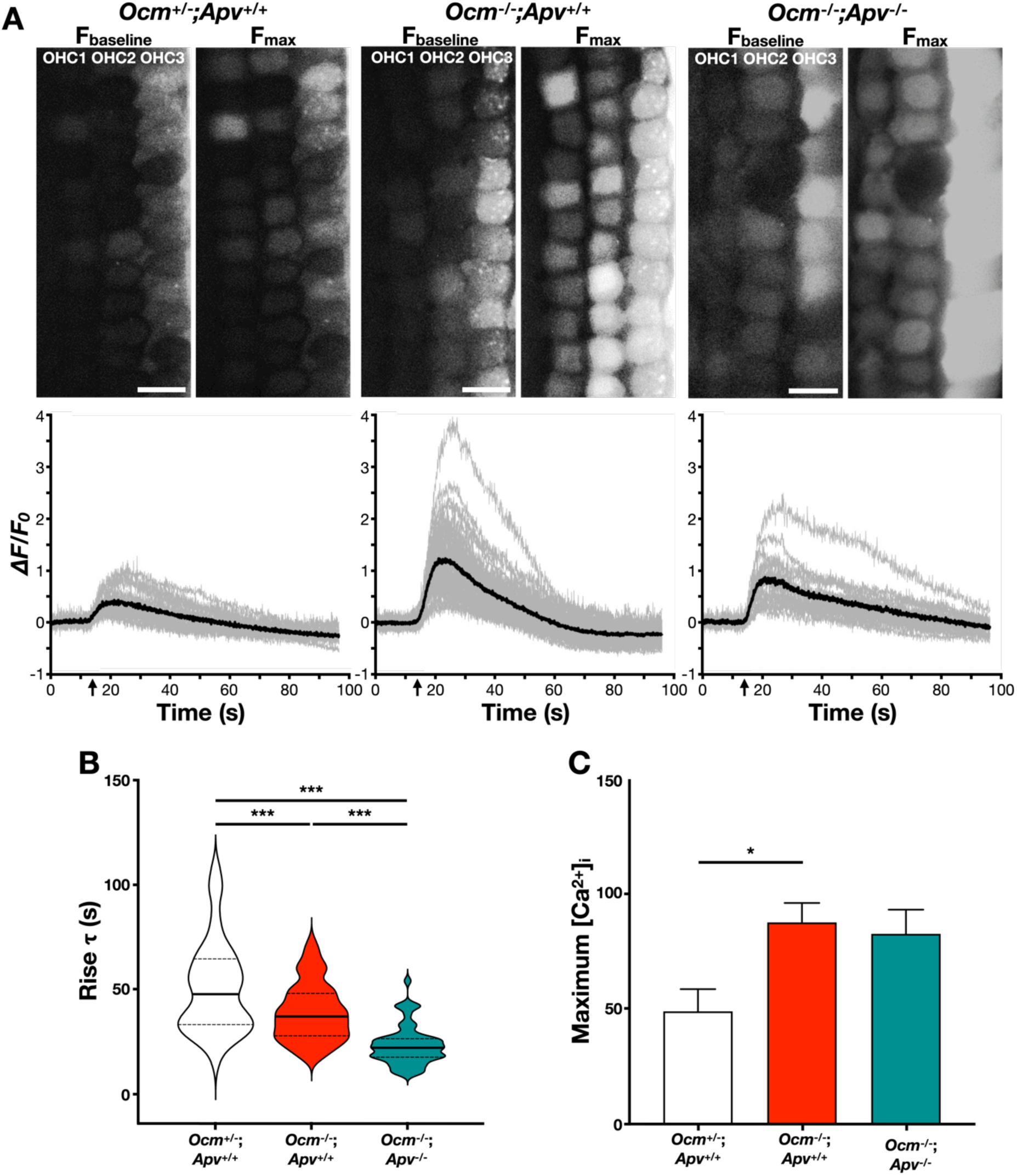
Calcium transients are altered in double *Ocm* and *Apv* knockout OHCs. **A)** Representative images depict Fbaseline (left: before stimulation with KCl) and Fmax (right: max change in fluorescence). Scale bar = 10 μm. Below *ΔF/F0* is plotted for each genotype. Grey lines are *ΔF/F0* measurements from individual OHCs. Solid black line represents the mean *ΔF/F0* for that preparation. **B)** Violin plots of rise τ for *Ocm*^+/-^;*Apv*^+/+^ (white), *Ocm*^-/-^;*Apv*^+/+^ (red), *Ocm*^-/-^;*Apv*^-/-^ (teal). n=3 animals per genotype. Solid lines represent medians. Dotted lines represent quartiles. Significant differences (*one-way ANOVA*) are denoted as follows: *\*\*\*p<0.001*). **C)** Fura-2 was used to estimate the [Ca^2+^]i levels (in nM) of *Ocm*^+/-^;*Apv*^+/+^ (black, n=38 OHCs) and *Ocm*^-/-^;*Apv*^+/+^ (red, n=36 OHCs), and *Ocm*^-/-^;*Apv*^-/-^ (teal, n=33 OHCs). Significance (*one-way ANOVA*) is denoted as *\*p=0.022*. n≥3 for each genotype.

According to the experiments described above, loss of both *Ocm* and *Apv* results in altered rise-time constants in P3 OHCs compared to the loss of *Ocm* alone. In order to quantify the levels of free intracellular Ca^2+^, we performed a similar set of experiments using the ratiometric Ca^2+^ indicator, Fura-2. First, we determined the double log calibration plot of Fura-2 (**Figure S13**). The Ca^2+^ response of the indicator was linear with an apparent K_d_ of 257.15 nM. Second, cochlear spirals from P3 mice were harvested, incubated with Fura-2, and depolarized via KCl superfusion (see Materials and Methods). Finally, the average maximum [Ca^2+^]_i_ was calculated for each genotype. Maximum [Ca^2+^]_i_ from *Ocm*^-/-^*;Apv*^+/+^ and *Ocm*^-/-^*;Apv*^-/-^ were virtually identical. However, the maximum [Ca^2+^]_i_ of *Ocm*^*+/-*^*;Apv*^*+/+*^ OHCs was almost 2-fold lower in comparison (**Figure 5C**) (*p=0.022, one-way ANOVA*). Taken together, we conclude that in P3 OHCs, deletion of *Ocm* results in an increase in free cytosolic Ca^2+^. Further, while deletion of *Apv* alters the kinetics of Ca^2+^ transients induced in *Ocm*^-/-^ OHCs, the total amount of freely available cytosolic Ca^2+^ is unchanged by additional deletion of *Apv*.

### 3.6. OCM effectively buffers Ca^2+^ in cultured cells

To investigate the general buffering capacities of OCM, APV, and sorcin, we compared changes in Ca^2+^ signaling in HEK293T cells expressing the above fluorescently tagged proteins. HEK293T cells endogenously express sorcin, little or no APV, and no endogenous OCM (Human Protein Atlas). We confirmed expression of *Ocm*-transfected HEK293T cells via RT-PCR (**Figure 6A**). First, we assessed the ability of OCM to buffer Ca^2+^ by inducing rises in intracellular Ca^2+^ waves with ATP, which stimulates release of Ca^2+^ from endoplasmic reticulum (ER) [41]. When stimulated with ATP, untransfected cells demonstrated a robust response compared to OCM expressing cells (**Figure 6B**). The ATP responses were abolished following depletion of intracellular Ca^2+^ stores by cyclopiazonic acid (CPA), consistent with previous reports [42], and confirm that the increases in cytosolic Ca^2+^ were from the ER.

**Figure 6.**
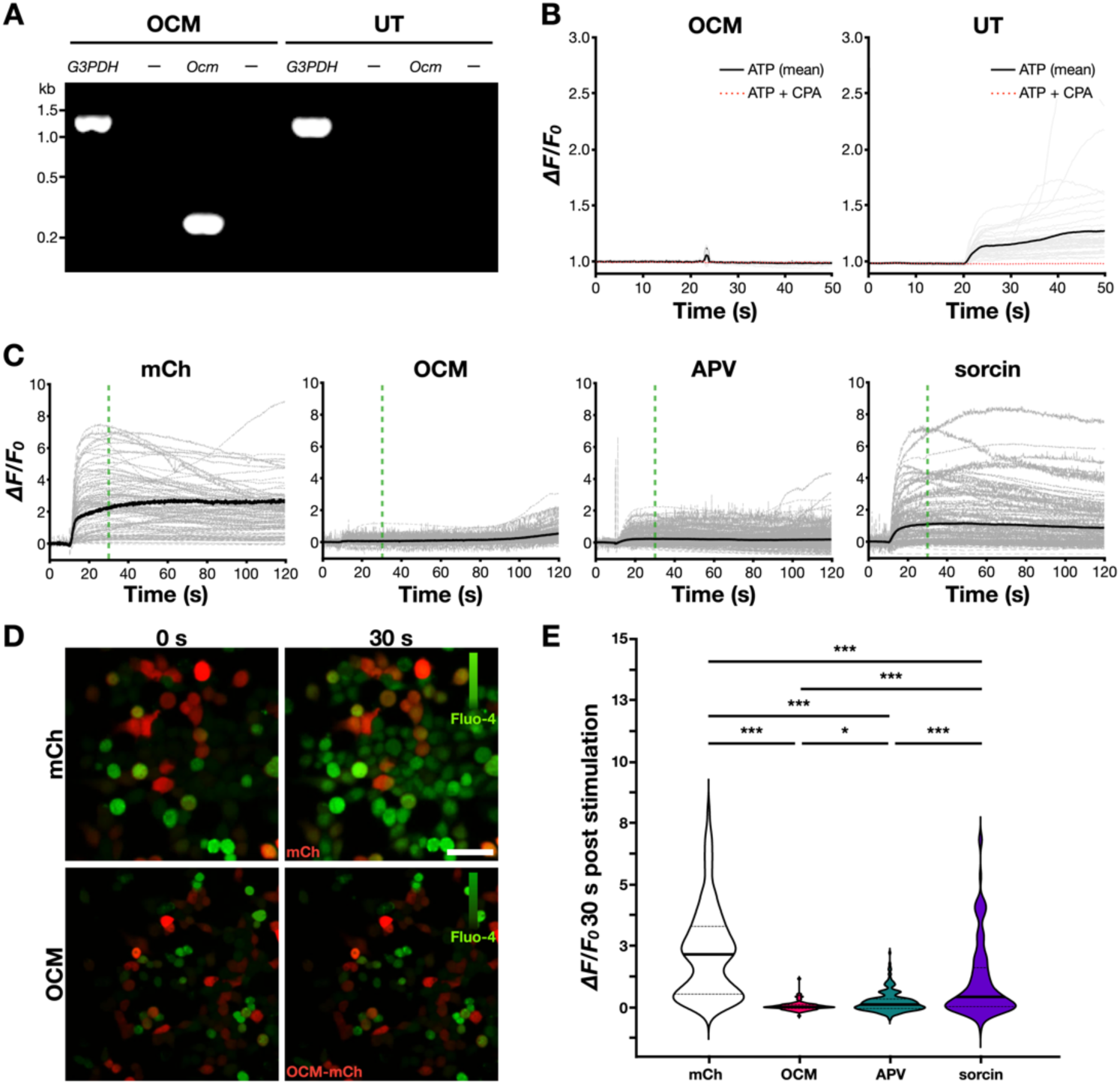
OCM influences calcium transient dynamics in transfected HEK293T cells. HEK293T cells were transiently transfected with fluorescently tagged CaBPs. **A)** *Ocm* mRNA is not endogenously detected in HEK293T cells. *G3PDH* and *Ocm* cDNA generated via RT-PCR from cell lysates of transfected (*Ocm*-CFP) and untransfected HEK293T cells. Wells labeled “-” are no transcript controls. **B)** HEK293T cells expressing fluorescently tagged OCM or control (untransfected) were incubated with Fluo-4 and stimulated with ATP. Changes in fluorescence of single cells are shown in grey. The solid black line indicates mean *ΔF/F0* for that group. Dotted red lines represent control experiments including CPA, a SERCA inhibitor. **C)** Representative plots of *ΔF/F0* upon stimulation with ionomycin. Changes in fluorescence of single cells are shown in grey. Solid black line indicates mean *ΔF/F0* for that group. Dashed green line marks 30 s post stimulation. **D)** Fluorescent proteins (red): mCh-only, *Ocm*-mCh, *Apv*-mCh (not shown), *Sri*-mCh (not shown) were incubated with Fluo-4 (green) and stimulated with ionomycin. Shown are still images before (0 s) and 30 s post-stimulation with ionomycin. Scale bar = 50 μm. **E)** Violin plots of *ΔF/F0* 30 seconds post stimulation with ionomycin for HEK293T cells positive for: mCh (white), OCM (pink), APV (teal), and sorcin (purple). Solid lines represent medians. Dashed lines represent quartiles. n≥100 cells per group. Significance (*one-way ANOVA*) denoted as: \**p=0.02* and \*\*\**p<0.001*

Next, we assessed the ability of OCM to buffer Ca^2+^ in HEK293T cells by application of ionomycin, an ionophore that increases cytosolic Ca^2+^ levels from exogenous sources [43, 44]. mCherry (mCh)-tagged OCM, APV and sorcin plasmids were separately transfected into HEK293T cells. A flexible linker was inserted in between sequences encoding the aforementioned CaBPs and the fluorescent tag (mCh) to more closely mimic the inherent binding kinetics of the proteins (see **Figure S01** for plasmid maps and protein configurations). Cells expressing the control plasmid (mCh) displayed rapid and highly variable changes in fluorescence in response to stimulation with ionomycin that peaked between 20 and 30 s (**Figure 6C**,**D**). At 30 seconds post stimulation with ionomycin, mCh-expressing cells had a mean *ΔF/F*_*0*_ of 2.25±1.82 (**Figure 6E**). In contrast, we observed a very slow, gradual *ΔF/F*_*0*_ from cells expressing *Ocm*-mCh, which, at 30 seconds post stimulation had a mean *ΔF/F*_*0*_ of 0.05±0.19.

Ionomycin-induced responses of HEK293T cells transfected with APV (0.21±0.40) and sorcin (1.08±1.54) showed significant decreases in *ΔF/F*_*0*_ at 30 s post-stimulation compared to control (*p<0.0001* for both comparisons, *one-way ANOVA*). (**Figure 6E**). Still, OCM expressing cells had the smallest variance and lowest *ΔF/F*_*0*_ at 30 s post-stimulation relative to both APV and sorcin expressing cells (*p=0.0173* and *p<0.0001*, respectively). Although all three CaBPs reduced changes in fluorescence after ionomycin-induced Ca^2+^ flux, OCM was more effective than either APV or sorcin at minimizing changes in [Ca^2+^]_i_. The results from our mammalian cell culture experiments align with experiments performed with OHCs. We conclude that OCM lowers the amount of freely available [Ca^2+^]_i_ and increases the Ca^2+^ buffering capacity in HEK293T.

## 4. Discussion

We present the first direct evidence that OCM plays a unique and important role as a Ca^2+^ buffer in developing OHCs. While OHCs from *Ocm*^*-/-*^ mice display normal membrane potentials and basolateral membrane currents, loss of OCM disrupts normal Ca^2+^ buffering dynamics. In P0 OHCs, when OCM expression is minimal to non-existent, deletion of *Ocm* causes no major change in Ca^2+^ transient dynamics. However, at P3, the onset of OCM protein expression, the rise time constants of Ca^2+^ signals are faster in *Ocm*^*-/-*^ compared to age-matched control *Ocm*^*+/-*^ mice. This indicates that during early stages of development, OCM plays a significant role in sculpting Ca^2+^ signaling in OHCs, which has been shown to promote the maturation of their biophysical characteristics and innervation [25, 37]. Deletion of both *Ocm* and *Apv* further changes OHC Ca^2+^ signaling, indicating that both OCM and APV shape Ca^2+^ dynamics in immature OHCs. Using a HEK293T cell culture model, we show that transiently expressed OCM rapidly buffers cytosolic Ca^2+^ more efficiently than either APV or sorcin. We conclude that OCM plays a significant role in the Ca^2+^ signaling network of immature OHCs.

In cochlear hair cells, EF-hand CaBPs are differentially regulated during the postnatal period. The present study found that *Ocm* mRNA expression turns on as early as P0, although at relatively low levels, and significantly increases by P3. This timeline is in accordance with RNA-seq analyses [45-49]. Our immunofluorescent analyses fall in line with previous studies, concluding that OCM protein expression increases during early postnatal OHC development, while APV expression decreases [12, 14]. Sorcin (soluble resistance-related calcium-binding protein) expression follows the same trend as APV. In control OHCs, we see a decrease in sorcin expression from P0 to P3. Recent studies show that sorcin is highly expressed in OHCs [17]. Sorcin is one of the most expressed Ca^2+^-binding proteins in the brain and cardiac tissues [50]. It is a key protein in the endoplasmic reticulum Ca^2+^-dependent cascades. In cardiac myocytes, sorcin regulates Ca^2+^ levels in the subsurface cisternae and inhibits Ca^2+^-induced Ca^2+^-release [16, 17]. Sorcin likely plays a similar role in OHCs; however, its exact mechanism has yet to be defined.

Earlier studies suggest that genetic deletion of one EF-hand CaBP does not typically lead to the upregulation of other CaBPs [51]. However there are some exceptions to this rule, such as the upregulation of OCM in the brain of *Apv*^*-/-*^ mice [38, 39]. Our investigations in the cochlea suggest that the presence or absence of OCM does alter the expression of other EF-hand CaBPs at early developmental stages. Protein expression of APV and sorcin is altered in cochleae of *Ocm*^*-/-*^ mice at both P0 and P3. However, the patterns in mRNA expression do not match the patterns observed in protein expression and the immunofluorescent changes are found in both OHCs and IHCs. At P3 the *Sri* mRNA expression decreases in the *Ocm* knockout whereas the sorcin-ir is more intense in *Ocm* knockout hair cells. There are multiple potential explanations for this. One possibility is that the lack of *Ocm* in P3 hair cells causes *Sri* expression to be differentially regulated in hair cells by some unknown post-transcriptional mechanism. Another possibility is that the mRNA expression studies are using whole tissues that include much more than just hair cells. In our qRT-PCR analysis, cochlear spirals are pooled, which include a diverse set of cells, all of which express *Sri* at high levels during development [48]. One benefit of immunofluorescent analysis is the ability to observe immunoreactivity in OHCs and IHCs, specifically. Although *Ocm* is preferentially expressed in OHCs, there is a developmentally transient and minimal expression of *Ocm* in IHCs [12, 17]. Here, we show that deletion of Ocm causes changes in sorcin immunofluorescence both in OHCs and IHCs. Thus, the presence of *Ocm* in IHCs, although relatively low, may contribute to the developmental expression of other CaBPs.

The dynamic changes in gene expression of OCM and APV during postnatal development suggest that these CaBPs may contribute to shaping the Ca^2+^ signaling network of OHCs in different ways. Here, the contribution of OCM in maintaining OHC Ca^2+^ homeostasis is highlighted in two ways: 1) At P0, before the onset of OCM expression, genetic deletion of *Ocm* has no effect on Ca^2+^ transients and 2) genetic deletion of *Ocm* at P3 causes considerable changes in Ca^2+^ rise-time kinetics. These changes in Ca^2+^ signaling cannot be attributed to differences in biophysical profiles. Regardless of any attempted compensational mechanisms, loss of *Ocm* leads to dysregulated Ca^2+^ signaling in OHCs. The present study suggests that APV also contributes somewhat to shaping Ca^2+^ transients in OHCs at P3. In the absence of OCM, the additional loss of APV leads to a further decrease in OHC rise-time kinetics. However, in *Ocm*^*-/-*^ OHCs, loss of *Apv* does not appear to contribute to the total maximum [Ca^2+^]_i_ measured in P3 OHCs. This discrepancy could reflect differences in Ca^2+^ binding kinetics between OCM and APV. APV is considered a slow-onset Ca^2+^ buffer [51], while OCM has not yet been defined as a slow-or fast-onset buffer.

Although OCM and APV share many similarities such as their sequence identity, molecular weights, and tertiary structures [10], at least in HEK293T cells, the two proteins demonstrate differences in Ca^2+^ buffering capacity. OCM and APV are distinguished by their metal ion binding properties [21, 35, 52-54]. Rapid Ca^2+^ buffering by OCM is supported by our mammalian cell culture model. We observe a slow, gradual *ΔF/F*_*0*_ in OCM expressing cells following treatment with ionomycin or ATP. While ionomycin causes increases in cytosolic Ca^2+^ levels from exogenous sources [43, 44], ATP increases cytosolic Ca^2+^ through an intracellular mechanism. ATP induces Ca^2+^ transients in HEK293T cells via P2Y receptors in an inositol triphosphate (IP_3_)–dependent manner [41, 55-60]. Pre-treatment with cyclopiazonic acid (CPA), a SERCA inhibitor, abolishes the change in fluorescence of the Ca^2+^-indicator dye seen with ATP-treatment, confirming that the rise in cytoplasmic [Ca^2+^]_i_ levels is caused by release of Ca^2+^ from ER stores [31]. OCM effectively buffers cytosolic Ca^2+^ levels, irrespective of the source, that is, stored vs. exogenous. Further, OCM buffers Ca^2+^ more effectively than either APV or sorcin.

While loss of *Ocm* alters *Apv* and *Sri* expression during development, these changes are ineffective in maintaining Ca^2+^ homeostasis over time, since young adult mice show accelerated age-related hearing loss [10, 18]. Other components of the Ca^2+^ signaling network, such as Ca^2+^ channels, development of the subsurface cisternae, localization of mitochondria, etc. may also be influenced by loss of *Ocm*. It would be worthwhile to explore these mechanisms and determine whether they are consequences of *Ocm* deletion that further exacerbate abnormal Ca^2+^ signaling or compensatory mechanisms that attempt to restore Ca^2+^ homeostasis. The dysregulated Ca^2+^ signaling we observe in neonatal *Ocm* mutant OHCs could ultimately lead to Ca^2+^ overload in adult OHCs, which may be linked to the progressive early-onset hearing loss phenotype observed in young adult mice [18]. We conclude that OCM plays a distinct and important role in the development of Ca^2+^ signaling in OHCs.

## Author Agreement

All authors have read and approved the final version.

## Author Contributions

DS and WM designed, planned and supervised the research. KM and YY contributed equally to experimental design and implementation of the work. DS, WM, FC, YY, and SD performed Ca^2+^ transient experiments. FC, JJ, and WM performed electrophysiology experiments. YY, FJ, and JC performed qRT-PCR and immunofluorescence. KM and LC performed cloning and cell culture. LC, YY, and AH bred and maintained mouse colonies. LC performed hearing tests. YY, FC, JJ, FJ, WM, and DS analyzed the data. FC, YY and DS wrote and implemented code for data analysis. KM, YY, and DS wrote the paper. All authors contributed to the editing of the paper.

## CRediT authorship contribution statement

**Kaitlin Murtha:** Investigation, Writing – Original Draft, Writing – Review & Editing, Visualization. **Yang Yang:** Methodology, Software, Validation, Formal analysis, Investigation, Writing – Review & Editing, Visualization. **Federico Ceriani:** Methodology, Validation, Formal Analysis, Investigation, Writing – Review & Editing. **Jin-Yi Jeng:** Methodology, Validation, Formal Analysis, Investigation, Writing – Review & Editing. **Leslie Climer:** Conceptualization, Investigation, Writing – Review & Editing, Supervision, Project administration. **Forrest Jones:** Investigation, Writing – Review & Editing. **Jack Charles:** Investigation, Writing – Review & Editing. **Sal Davana:** Investigation, Writing – Review & Editing. **Aubrey Hornak:** Project administration, Investigation, Writing – Review & Editing. **Walter Marcotti:** Conceptualization, Methodology, Software, Resources, Writing – Review & Editing, Supervision, Project administration, Funding acquisition. **Dwayne Simmons:** Conceptualization, Methodology, Formal analysis, Investigation, Resources, Data Curation, Writing – Original Draft, Writing – Review & Editing, Visualization, Supervision, Project administration, Funding acquisition.

## Acknowledgements

This research was supported by National Institute on Deafness and Other Communication Disorders Grants DC013304 and DC018935 (DDS), 2015-2016 Fulbright U.S. Scholar Award 5403 (DDS), American Hearing Research Foundation grant, the BBSRC BB/T004991/1 (WM).

## Supplemental Figure Legends

**Figure S01**. Plasmid and Gene Sequences. Rat *Ocm, Apv, Sri* nucleotide sequences, p*Ocm*-CFP, p*Ocm*-mCh, p*Apv*-mCh, p*Sri*-mCh, pmCh plasmid nucleotide sequences and plasmid maps.

**Figure S02**. CFP and Fluo-4 excitation filters for Ca^2+^ transients in mammalian cells. Excitation (dotted lines, no fill) and emission (color filled) plots are shown for both fluorophores. Filters are shown as highlighted rectangles. Visual created using Fluorescence SpectraViewer (ThermoFisher).

**Figure S03**. Primer Sequences. Table of primer sequences used for qRT-PCR and PCR. Text Figure.

**Figure S04**. qRT-PCR Data and Calculations. Spreadsheets containing raw Cq values, 2-(ΔΔCq) values, and calculations for all qRT-PCR experiments.

**Figure S05**. P0 *Ocm*^*+/-*^ local KCl application Calcium Transient. Video (.avi) file of Fluo-4 incubated P0 *Ocm*^+/-^ spiral stimulated with KCl. 19.70 s.

**Figure S06**. P0 *Ocm*^*-/-*^ local KCl application Calcium Transient. Video (.avi) file of Fluo-4 incubated P0 *Ocm*^-/-^ spiral stimulated with KCl. 19.70 s.

**Figure S07**. P3 *Ocm*^*+/-*^ local KCl application Calcium Transient. Video (.avi) file of Fluo-4 incubated P3 *Ocm*^+/-^ spiral stimulated with KCl. 06.50 s.

**Figure S08**. P3 *Ocm*^*-/-*^ local KCl application Calcium Transient. Video (.avi) file of Fluo-4 incubated P3 *Ocm*^-/-^ spiral stimulated with KCl. 06.50 s.

**Figure S09**. DPOAE threshold measurements (dB SPL) of 1 month-old C57Bl/6 *Ocm*^*+/+*^*;Apv*^*+/+*^ (n=7, black squares), *Ocm*^*-/-*^*;Apv*^*+/+*^ (n=9, red filled circles) and *Ocm*^*-/-*^*;Apv*^*-/-*^ (n=10, red outlined circles) C57Bl/6 mice. Mean and SEM are plotted. **significant difference at 32 kHz between *Ocm*^*+/+*^*;Apv*^*+/+*^ and *Ocm*^*-/-*^*;Apv*^*+/+*^ (*p=0.0021*, Tukey’s multiple comparisons).

**Figure S10**. P3 *Ocm*^*+/-*^*;Apv*^*+/+*^ KCl superfusion Calcium Transient. Video file (.avi) of Fluo-4 incubated dissected organ of Corti spiral stimulated via KCl superfusion. 233.33 s.

**Figure S11**. P3 *Ocm*^*+/-*^*;Apv*^*-/-*^ KCl superfusion Calcium Transient. Video file (.avi) of Fluo-4 incubated dissected organ of Corti spiral stimulated via KCl superfusion. 200.00 s.

**Figure S12**. P3 *Ocm*^*-/-*^*;Apv*^*-/-*^ KCl superfusion Calcium Transient. Video file (.avi) of Fluo-4 incubated dissected organ of Corti spiral stimulated via KCl superfusion. 200.00 s.

**Figure S13**. Fura-2 Calibration Curve.

